# Differences in transcription initiation directionality underlie distinctions between plants and animals in chromatin modification patterns at genes and cis-regulatory elements

**DOI:** 10.1101/2023.11.03.565513

**Authors:** Brianna D. Silver, Courtney G. Willett, Kelsey A. Maher, Dongxue Wang, Roger B. Deal

## Abstract

Transcriptional initiation is among the first regulated steps controlling eukaryotic gene expression. High-throughput profiling of fungal and animal genomes has revealed that RNA Polymerase II (Pol II) often initiates transcription in both directions at the promoter transcription start site (TSS), but generally only elongates productively into the gene body. Additionally, Pol II can initiate transcription in both directions at cis-regulatory elements (CREs) such as enhancers. These bidirectional Pol II initiation events can be observed directly with methods that capture nascent transcripts, and they are also revealed indirectly by the presence of transcription-associated histone modifications on both sides of the TSS or CRE. Previous studies have shown that nascent RNAs and transcription-associated histone modifications in the model plant *Arabidopsis thaliana* accumulate mainly in the gene body, suggesting that transcription does not initiate widely in the upstream direction from genes in this plant. We compared transcription-associated histone modifications and nascent transcripts at both TSSs and CREs in *Arabidopsis thaliana, Drosophila melanogaster*, and *Homo sapiens*. Our results provide evidence for mostly unidirectional Pol II initiation at both promoters and gene-proximal CREs of *Arabidopsis thaliana*, whereas bidirectional transcription initiation is observed widely at promoters in both *Drosophila melanogaster* and *Homo sapiens*, as well as CREs in *Drosophila*. Furthermore, the distribution of transcription-associated histone modifications around TSSs in the *Oryza sativa* (rice) and *Glycine max* (soybean) genomes suggests that unidirectional transcription initiation is the norm in these genomes as well. These results suggest that there are fundamental differences in transcriptional initiation directionality between flowering plant and metazoan genomes, which are manifested as distinct patterns of chromatin modifications around RNA polymerase initiation sites.

## INTRODUCTION

All organisms must control gene expression in a manner that is both cell type-specific and adaptive to changing cues. As such, transcription is a highly dynamic and regulated process, with many conserved mechanisms across eukaryotes. However, the nuanced differences in transcriptional regulation between eukaryotic kingdoms and even individual species are still being uncovered.

In an effort to further understand the relationships between transcription and chromatin, previous studies have analyzed correlations between transcription and histone post-translational modification (PTM) patterns. This led to the discovery of relationships between histone PTMs and different genomic regions, wherein unique PTM “chromatin signatures” can reflect whether a region is actively transcribed, poised, or constitutively silenced (Strahl and Allis 2000; Sims *et al*. 2003). As a pertinent example, H3K4me3 and H3K4me1 are an indirect result of RNA polymerase II (Pol II) activity. The Pol II-associated histone methyltransferase SET1 is responsible for the majority of H3K4 methylation in active genes in many eukaryotes, and its activity within the COMPASS complex leads to the distinct patterning of increased trimethylation at the TSS, dimethylation across the gene body, and monomethylation towards the 3’ end (Liu *et al*. 2005; Pokholok *et al*. 2005; Barski *et al*. 2007; Soares *et al*. 2017). In short, observing patterns of PTMs in conjunction with nascent transcript data such as Global Run-On Sequencing (GRO-seq)(Core *et al*. 2008), Native Elongating Transcript sequencing (NET-seq) (Mayer *et al*. 2015) or Nascent 5-EU-labeled RNA sequencing (Neu-seq) (Szabo *et al*. 2020) can be a powerful tool for furthering our understanding of transcriptional activity and directionality.

It has been observed that in many animal genomes the distribution of histone PTMs indicative of active transcription is bimodal around the transcription start site (TSS), suggesting that transcriptional initiation is bidirectional in these eukaryotes (Heintzman *et al*. 2007). Indeed, it has been demonstrated that Pol II often initiates in both directions at a given TSS in yeast and animals, regardless of the presence or absence of a gene on the opposite strand (Core *et al*. 2008; Neil *et al*. 2009; Seila *et al*. 2009; Xu *et al*. 2009). In contrast, the plant *Arabidopsis thaliana* shows these same histone PTMs flanking just one side of the TSS, suggesting a unidirectional transcriptional mechanism (Roudier *et al*. 2011). This curious observation, which may indicate fundamental differences in the mechanism of Pol II initiation between plants and animals, prompted us to investigate transcriptional patterns at both TSSs and cis-regulatory elements (CREs) by direct comparisons of diverse species.

Enhancers are CREs found in many organisms, including eukaryotes (Schwaiger *et al*. 2014; Villar *et al*. 2015; Zhu *et al*. 2015; Weber *et al*. 2016), bacteria (Xu and Hoover 2001) and viruses (Berg *et al*. 1984). On a molecular scale, enhancer sequences are comprised of a modular collection of transcription factor binding motifs which act as an assembly platform for *trans*-acting factors (Lee and Young 2000; Spitz and Furlong 2012). Sequence-specific transcription factors (TFs), general TFs, and co-factors associate with the enhancer and in turn recruit larger molecular machinery, including the Mediator complex, Pol II, nucleosome remodelers, and histone modifying proteins such as CPB/p300 to initiate transcription at the promoter (Vernimmen and Bickmore 2015). Given that the binding of sequence-specific DNA binding proteins at enhancers displaces nucleosomes, these sites tend to be hypersensitive to nuclease digestion and can thus be identified at large through assays such as DNase-seq (Boyle *et al*. 2008) (Song and Crawford 2010) and ATAC-seq (Buenrostro *et al*. 2013). Additionally, studies in animal systems have identified characteristic histone PTMs associated with the nucleosomes that flank these CREs (Wang *et al*. 2008; Hawkins *et al*. 2010; Ernst *et al*. 2011; Zentner *et al*. 2011; Bonn *et al*. 2012). The set of histone PTMs associated with CREs in a variety of cell types and species include H3K4me1 and H3K4me3, which are deposited co-transcriptionally, in combination with H3K27ac or H3K27me3, depending on the activity state of the enhancer. Recent evidence indicates that Pol II initiates at enhancer elements to generate enhancer RNAs (eRNAs) and this initiation, like at TSSs, is frequently bidirectional in animals (Wang *et al*. 2011; Pan *et al*. 2021).

In this study, we integrated chromatin accessibility, ChIP-seq, and nascent transcript data from *Arabidopsis thaliana, Homo sapiens*, and *Drosophila melanogaster* to explore interspecies differences in chromatin modifications and transcriptional directionality. We first show evidence for mostly unidirectional transcription at TSSs in Arabidopsis and bidirectional transcription at those of human and Drosophila. We then examined putative CREs, defined as nuclease hypersensitive intergenic sites. Using the conserved set of enhancer histone PTMs H3K27ac, H3K27me3, H3K4me1, and H3K4me3, in conjunction with chromatin accessibility and nascent transcript data, we also find differences in transcriptional directionality at CREs between animal and plant genomes. While Drosophila shows frequent bimodal production of eRNAs and bimodal deposition of PTMs at accessible CREs, Arabidopsis shows mainly unidirectional nascent RNA production, with PTMs flanking only the transcribed side of the accessible chromatin region. Furthermore, analysis of ChIP-seq data from *Oryza sativa* (rice) and *Glycine max* (soybean) suggests that this unimodal histone PTM pattern is not specific to *Arabidopsis* and may represent a fundamental difference in Pol II initiation processes between the plant and animal kingdoms. Taken together, our analyses provide additional insight into the transcriptional dynamics of plants and suggest that differences in transcriptional directionality underlie the disparities observed in chromatin modification patterns between plant and animal epigenomes.

## RESULTS

### PROMOTER TRANSCRIPTION IS BIDIRECTIONAL IN ANIMAL MODELS AND PREFERENTIALLY UNIDIRECTIONAL IN ARABIDOPSIS

To address transcriptional directionality at the TSSs of protein coding genes, we combined nuclease hypersensitivity, ChIP-seq, and GRO-seq data from *Homo sapiens* (human), *Drosophila melanogaster*, and *Arabidopsis thaliana* (Supplementary Table 1). Publicly available data were used for human myeloid cells (all data), Drosophila S2 cells, (all data) and Arabidopsis root epidermal non-hair cells (ATAC-seq and ChIP-seq) and Arabidopsis seedlings (nascent transcript data). ChIP-seq data from Arabidopsis were generated from root epidermal non-hair cells in this study (Supplementary Tables 2 and 3). As much as possible, we attempted to analyze data from single cell types in order to minimize signal interference from different cell types. For the human analysis, this meant that DNase-seq and ChIP-seq are from CD34+ myeloid progenitor cells (Bernstein *et al*. 2010), while the GRO-seq analysis was from CD34+ myeloid progenitor cells cultured for 14 days and analyzed before terminal differentiation into erythrocytes (Gao *et al*. 2017). For Arabidopsis, ATAC-seq (Maher *et al*. 2018) and ChIP-seq were from root epidermal non-hair cells but nascent transcriptome data were from seedlings (Hetzel *et al*. 2016; Zhu *et al*. 2018; Szabo *et al*. 2020). Finally, Drosophila DNase-seq, ChIP-seq, and GRO-seq all came from S2 cells (Herz *et al*. 2012) (Core *et al*. 2012). This means that at a minimum, accessibility and histone PTM data are from the same cell type in each organism.

Gene-centric metaplots of the average ChIP-seq signal for H3K27ac, H3K27me3, H3K4me1, and H3K4me3 enrichment and chromatin accessibility across gene bodies in each of the three species are shown in **Figure 1A**. At this global scale, broad similarities are apparent in the pattern of chromatin accessibility relative to gene bodies. The region of maximum accessibility is restricted to a narrow peak 100-250 bp directly upstream of the TSS. Despite this fundamental similarity among organisms, a striking distinction emerges when the enrichments of H3K27ac, H3K27me3, H3K4me1, and H3K4me3 are considered. The signal for these four histone modifications is clustered in a distinct bimodal pattern around the transcription start site (TSS) for both the human and Drosophila metaplots, with clear signals upstream and downstream of the TSS **(Figure 1A)**. This pattern of enrichment is attributed to the bidirectional nature of animal promoters and their proclivity to produce transcripts from a single TSS in both the sense and antisense directions (Trinklein *et al*. 2004; Kim *et al*. 2005; Barski *et al*. 2007; Guenther *et al*. 2007; Core *et al*. 2008). The elongating form of Pol II acts as a binding platform for histone modifying complexes, such as MLL3 and MLL4 in mammals, which deposit H3K4 methylation on the underlying histones successively through multiple rounds of elongation (Kaikkonen *et al*. 2013). As such, the process of transcription itself is responsible for the surrounding deposition of this characteristic set of histone modifications (Seila *et al*. 2009). This process leads to the enrichment of histone PTMs both upstream and downstream of the accessible TSS region in animals.

**Figure 1:**
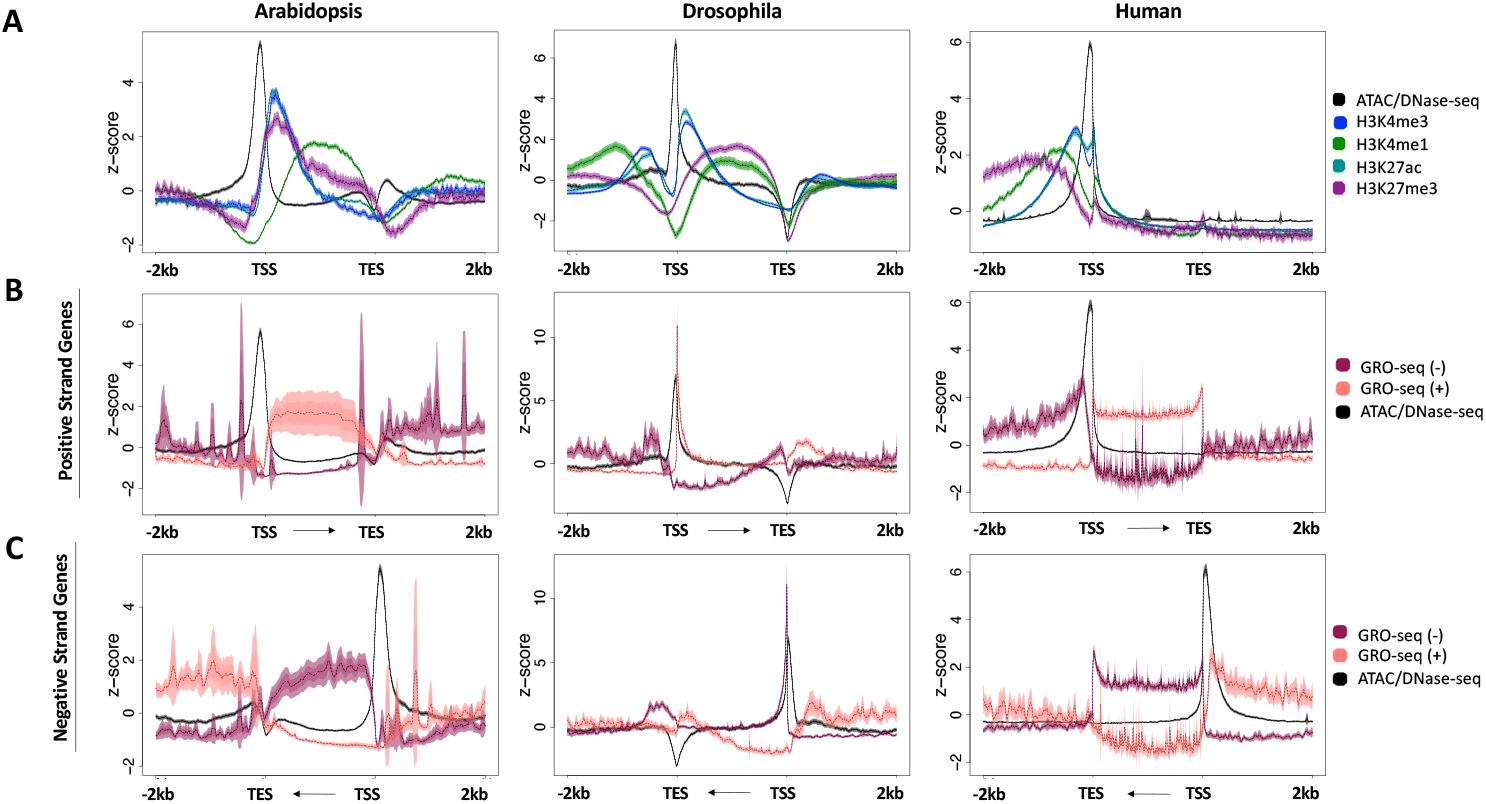
Histone modification enrichment and nascent transcripts across gene bodies in different organisms. A) Metaplots of average gene profiles of annotated Arabidopsis, Drosophila, and Human protein-coding genes. ChIP-seq signal for H3K27ac, H3K27me3, H3K4me1, and H3K4me3 are shown, along with chromatin accessibility data (ATAC-seq for Arabidopsis data; DNase-seq for Drosophila and Human data). B) Metaplots of the nascent transcriptional output (GRO-seq data) on Arabidopsis, Drosophila, and Human annotated positive strand and C) negative strand genes. Windows extend 2 kb upstream of the transcription start site (TSS) and 2 kb downstream of the transcript end site (TES). For each dataset, several statistical metrics are shown. Solid lines represent the mean signal intensity (z-score normalized); the inner, dark-shaded region represents the standard error; and the outer, light-shaded region represents the 95% confidence interval (CI) of signal intensity.

In contrast, a unique pattern is seen at the TSS of the Arabidopsis metaplot. The histone modification ChIP-seq signals are most abundant at the 5’ end of gene bodies, with the signal upstream of transcription start sites reduced to near background levels. Considering the mechanisms responsible for generating and maintaining the bimodal enrichment of histone modifications around animal TSSs, the absence of this pattern suggests that transcription in Arabidopsis may proceed nearly exclusively in the sense orientation, accounting for the sole downstream presence of histone marks. To examine this possibility more closely, we analyzed publicly available Global Run-On sequencing (GRO-seq) data from human, Drosophila, and Arabidopsis (Core *et al*. 2008; Hetzel *et al*. 2016; Gao *et al*. 2017). We separated all protein-coding genes across these genomes by their strandedness, plotting GRO-seq signal at all plus strand genes (**Figure 1B**) and minus strand genes (**Figure 1C**) separately. Within gene bodies, the Arabidopsis metaplots reveal that the directionality of the transcripts produced matches the directionality of the gene itself. In short, positive strand genes produce positive strand transcripts, while negative strand genes produce negative strand transcripts. Just upstream of the TSS in human and Drosophila genomes, transcripts running opposite of the genic direction are also produced, as is typical of bidirectional transcription at promoters (Kapranov *et al*. 2007; Core *et al*. 2008; Seila *et al*. 2008) and enhancers (Kim *et al*. 2010; Hah *et al*. 2013; Shlyueva *et al*. 2014). This lack of upstream signal in Arabidopsis indicates that transcription is strongly biased to be unidirectional in this organism, as originally observed by Hetzel and co-workers (Hetzel *et al*. 2016). This pattern of nascent transcripts matches that observed in the enrichment of histone PTM signals, and further supports that histone PTM enrichment reflects transcriptional output. These findings were further confirmed by analyzing two additional types of nascent transcript data at TSSs in Arabidopsis, NET-seq and 5-EU-RNA-seq (Zhu *et al*. 2018; Szabo *et al*. 2020) (Supplementary Figure 1A and B).

### INTERGENIC CREs ARE UNIDIRECTIONALLY FLANKED BY CHARACTERISTIC ENHANCER MARKS IN ARABIDOPSIS

We next sought to investigate whether the pattern of transcriptional directionality observed at TSSs persists at cis-regulatory elements (CREs) in Arabidopsis. CREs have been shown to preferentially reside in regions of hyperaccessible chromatin (Tsompana and Buck 2014; Jiang 2015), and accessible sites have thus been used as markers of putative regulatory elements, such as enhancers (Bell *et al*. 2011). To examine and compare gene-proximal CREs across species, we defined intergenic accessible regions (IARs) as nuclease hypersensitive sites that were outside of transcribed protein coding regions and were in the range of 100-2,000 bp away from a TSS to reduce signals from protein-coding gene TSSs. We focused on this window for two reasons. First, the majority of non-genic CREs in the *Arabidopsis*, rice, tomato, and *Medicago* genomes fall within 2 kb of the TSS (Maher *et al*. 2018) and many CREs have also been observed relatively close to the TSS in a variety of other angiosperm species, both monocots and dicots (Lu *et al*. 2019). Second, analysis of enhancer element positioning by STARR-seq in plants suggested that enhancers may preferentially function in the upstream orientation (Jores *et al*. 2020), thus the IARs might be enriched for bona fide enhancer elements. To assess whether RNA signals flanking IARs represent only stable transcripts or whether some represent transient RNAs, as would be expected for CRE-derived RNAs, we compared GRO-seq and steady-state RNA-seq in these regions (Supplementary Figure 2). We found that while GRO-seq and RNA-seq signal frequently overlap, there also appears to be a substantial proportion of sites that display GRO-seq signal but not RNA-seq signal, consistent with these representing unstable RNAs. For the sake of consistency across analyses, we also used this same window to select IARs when analyzing the Drosophila and human datasets.

We began by mapping four enhancer-associated histone modifications onto IARs. While regulatory elements are nucleosome-depleted regions where the frequent binding of *trans-*acting factors leaves the chromatin highly accessible, well-positioned nucleosomes tend to flank the boundaries of these regions, often carrying characteristic histone modifications (Schones *et al*. 2008; Henikoff *et al*. 2009; Jin *et al*. 2009). The metaplots of histone modification enrichment at IARs in all three species (**Figure 2A**) show symmetrical enrichment for H3K27ac, H3K4me1, and H3K4me3, which is in line with what has been previously reported in animal studies. Much like the pattern at gene bodies (**Figure 1**), H3K4me3 is enriched close to the accessible region, with H3K4me1 enrichment appearing more distally. These modifications are deposited during the process of transcription as the polymerase transitions from initiation to elongation (Kaikkonen *et al*. 2013), and are indicative of the production of RNAs surrounding the accessible chromatin region (**Figure 2A**). GRO-seq signal was also mapped over IARs (**Figure 2B**), revealing the presence of nascent RNAs arising from these regions in all three species.

**Figure 2:**
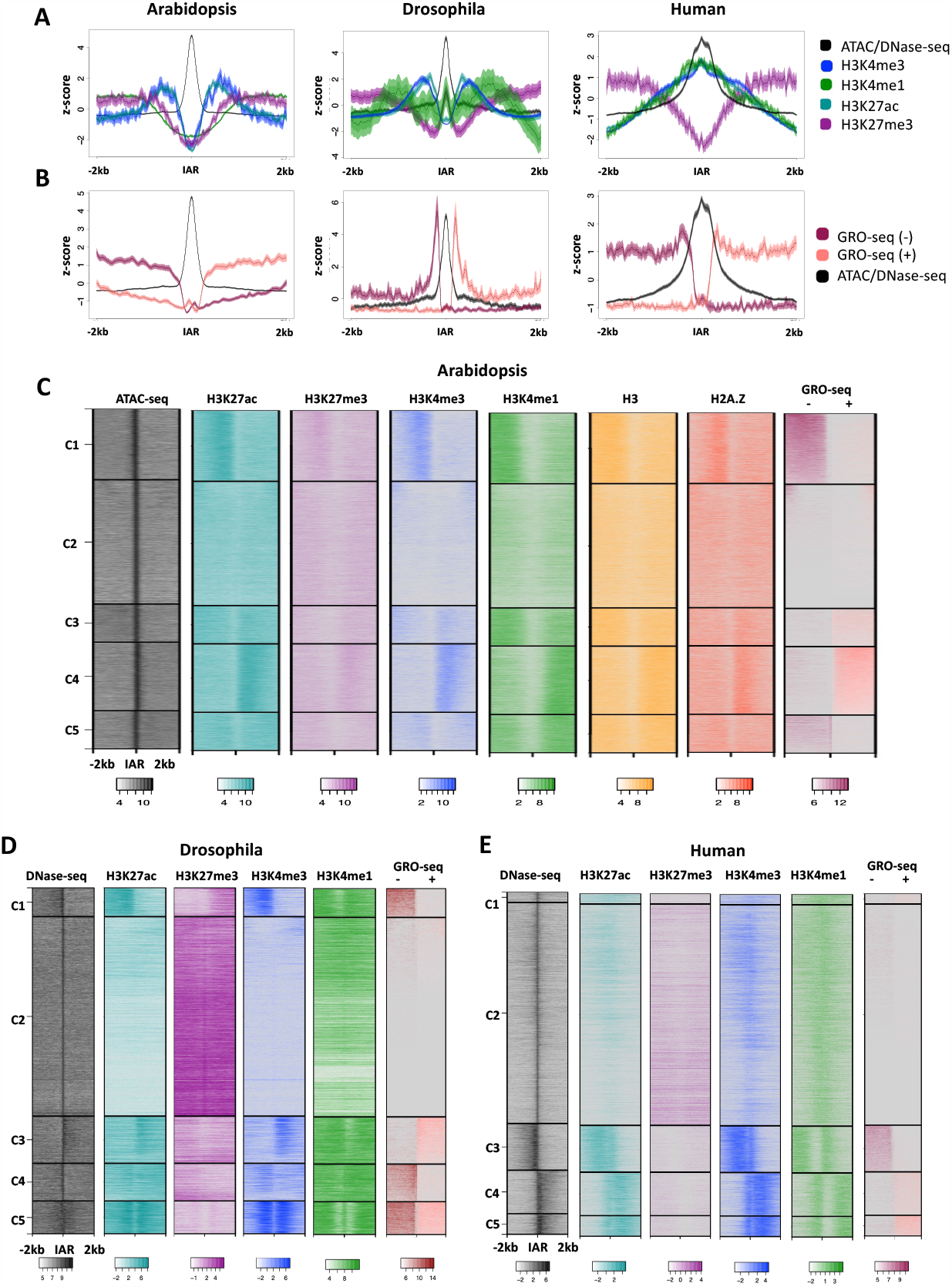
Histone modification enrichment and nascent transcript patterns around intergenic accessible chromatin regions (IARs). **A)** Metaplots of average histone modification and chromatin accessibility profiles at proximal intergenic accessible regions (IARs) in Arabidopsis, Drosophila, and Human. **B)** Metaplot of chromatin accessibility and nascent transcriptional output (GRO-seq) at IARs. In both A and B, solid lines represent the mean signal intensity (z-score normalized); the inner, dark-shaded region represents the standard error; and the outer, light-shaded region represents the 95% confidence interval (CI) of signal intensity. **C)** Heatmaps of average profiles of proximal intergenic accessible regions in Arabidopsis (19,962), **D)** Drosophila (7,702 sites), and **E)** Human (12,664 sites). Heatmaps were log2 transformed and divided into 5 k-means clusters. Windows extend 2 kb upstream and 2 kb downstream of each accessible site.

While metaplots are useful in displaying the average signal across a group of loci, heatmaps expand on these trends by showing the signal pattern at each unique locus. Intergenic accessible chromatin regions across the *Arabidopsis*, Drosophila, and human genomes are displayed in a heatmap, grouped into subpopulations via k-means clustering in **Figure 2C-E**. In addition to H3K27ac, H3K27me3, H3K4me1, H3K4me3, and chromatin accessibility data, we also generated ChIP-seq datasets for histones H3 and H2A.Z in the *Arabidopsis* root epidermal non-hair cell type, which are also displayed in **Figure 2C**. H2A.Z is a histone variant associated with the flanking regions of active enhancers, while histone H3 is a core component of histone octamers and a marker of nucleosome occupancy (Jin *et al*. 2009). High intensity signal can be seen in the center of the window in each ATAC-seq or DNase-seq heatmap, indicating a pronounced region of high chromatin accessibility. While there are clear differences between each of the species regarding the patterning of ChIP-seq and GRO-seq results, some universal patterns are observed. Generally, active transcription as indicated by the presence of GRO-seq signal overlaps with H3K4me3, H3K4me1, and H3K27ac. This pattern is consistent with the presence of active enhancers, which are producing nascent RNAs. In each case H3K27ac and H3K4me3 are generally enriched closest to the center of the IAR, followed by H3K4me1 (**Figure 2C-E**). As has been documented previously in eukaryotic genes, lysine 4 of histone 3 is predominantly trimethylated at the 5’ end of genes, with the modifications progressing to di- and monomethylation as transcriptional elongation proceeds (Shilatifard 2006; Li *et al*. 2007). Finally, the Arabidopsis H2A.Z and H3 data show enrichments corresponding to higher GRO-seq signal, suggesting that these nucleosomes are particularly well positioned, perhaps promoting transcription. (Schones *et al*. 2008; Henikoff *et al*. 2009; Jin *et al*. 2009).

Distinct from most studied eukaryotes, however, the Arabidopsis chromatin shows enrichment for both H3K27ac and H3K27me3 at the same loci **(Figure 2C, clusters 1 and 4)**. This is in contrast to the pattern shown in **Figures 2D** and **2E**, which show more exclusivity of H3K27ac and H3K27me3 in Drosophila and human, respectively. While these modifications are considered to be mutually exclusive in animal models, this simultaneous enrichment has been documented in previous Arabidopsis chromatin studies (Zhu *et al*. 2015). Whether this is due to the presence of nucleosomes that are dually enriched with the marks – bearing one H3 with lysine 27 methylated, the other H3 lysine 27 acetylated – or due to different chromatin states among the genome copies in polyploid cells, is not yet clear.

Regarding transcription at IARs, the GRO-seq data suggest a clear preference for initiation in only one direction in Arabidopsis (**Figure 2C, Clusters 1**,**4**). The same unidirectional transcription pattern observed in clusters C1 and C4 in Arabidopsis was also observed in publicly available NET-seq and 5-EU RNA-seq (Supplementary Figure 1C and D) (Zhu *et al*. 2018; Szabo *et al*. 2020). Consistent with this observation, eRNAs from Arabidopsis immunity-related CREs were also shown to be mostly unidirectional (Zhang *et al*. 2022).

In contrast, *Drosophila* IARs display enrichment patterns consistent with either bidirectional (C5) or unidirectional (C1,3,4) transcription (**Figure 2D**). Many clusters show dual enrichment of H3K4me3 and H3K4me1, indicative of transcription (Clusters 1,3,4,5). In addition, many of the loci within these clusters also show moderate H3K27ac signal, as is typical of ‘active’ enhancers (Creyghton *et al*. 2010; Rada-Iglesias *et al*. 2011; Bae and Lesch 2020). Other clusters show enrichment for H3K27me3/H3K4me1 characteristic of ‘poised/inactive’ enhancers (Cluster 2) (Creyghton *et al*. 2010; Rada-Iglesias *et al*. 2011).

Similar to *Arabidopsis*, the human heatmaps exclusively show a preference for unidirectionality **(Figure 2E)**. GRO-seq enrichment is only seen on one side of the IAR, as at least partly mirrored by ChIP-seq signal (Cluster 3,4,5). This was somewhat surprising, as most literature supports bidirectional transcription of eRNAs in human cells (Melgar *et al*. 2011; Hah *et al*. 2013). One possibility is that the human gene-proximal CREs examined here behave differently than those at large. Alternatively, it could be that the erythroid cell type we are analyzing does in fact preferentially produce eRNA transcripts in a unidirectional manner. Erythroblasts are the last stage before terminal differentiation, and during the final step to becoming a red blood cell, the nucleus is expelled. Just prior to this, the amount of RNA Polymerase II in the nucleus drops considerably, and it could be that with less overall transcriptional initiation at this stage, the production of bidirectionally transcribed eRNAs decreases (Larke *et al*. 2021; Murphy *et al*. 2021). This notion is consistent with reports that higher transcriptional activity correlates with increased production of bidirectional transcripts and eRNAs (Andersson *et al*. 2014).

Finally, closer investigation into the relationships between nascent transcripts and histone PTMs reveals some notable differences between human and *Arabidopsis*. In each unidirectional cluster (**Figure 2E** clusters C3, 4, 5) in human, there is enrichment of H3K4me3 and H3K4me1 on both the transcribed and the non-transcribed side of the IAR, which is most reminiscent of the Drosophila bidirectional cluster (**Figure 2D**, cluster C5), however, the signal does not appear to stretch as far across the 2kb window in humans. It is possible that the ChIP-seq pattern present is a remaining hallmark of previous bidirectional transcription that started to diminish with lower rates of transcriptional activity. This is supported by a recent study that concluded that terminal erythroid maturation is associated with a loss of histone marks indicative of transcriptional elongation, but without a corresponding increase in heterochromatin marks (Murphy *et al*. 2021).

In addition to examining transcriptional direction patterns qualitatively, we sought to quantify directionality. To calculate the bias of transcriptional signal across IARs observed in each genome, we calculated the average GRO-seq signal both upstream and downstream of each IAR in each species. Statistical significance of differences between the two directions at a given IAR was determined using the Wilcoxon rank sum test, a non-parametric version of the two-sample t-test (Lin *et al*. 2021) and a Bonferroni-corrected significance threshold was applied (Curtin and Schulz 1998) **(Figure 3)**. The apparent unidirectionality of Clusters C1 and C4 in Arabidopsis were supported by a statistically significant difference (p < 2.2e-16) between upstream and downstream signal (**Figure 3A)**. In contrast, Cluster C5 in Drosophila had an insignificant (p = 0.01167) difference between up and downstream signal, indicative of bidirectional enrichment of GRO-seq signal (**Figure 3B)**. As suggested by the qualitative analysis in Figure 2, human IARs preferentially showed unidirectional transcription, as indicated by significant differences in read density between the sides of each transcribed IAR (**Figure 3C**).

**Figure 3:**
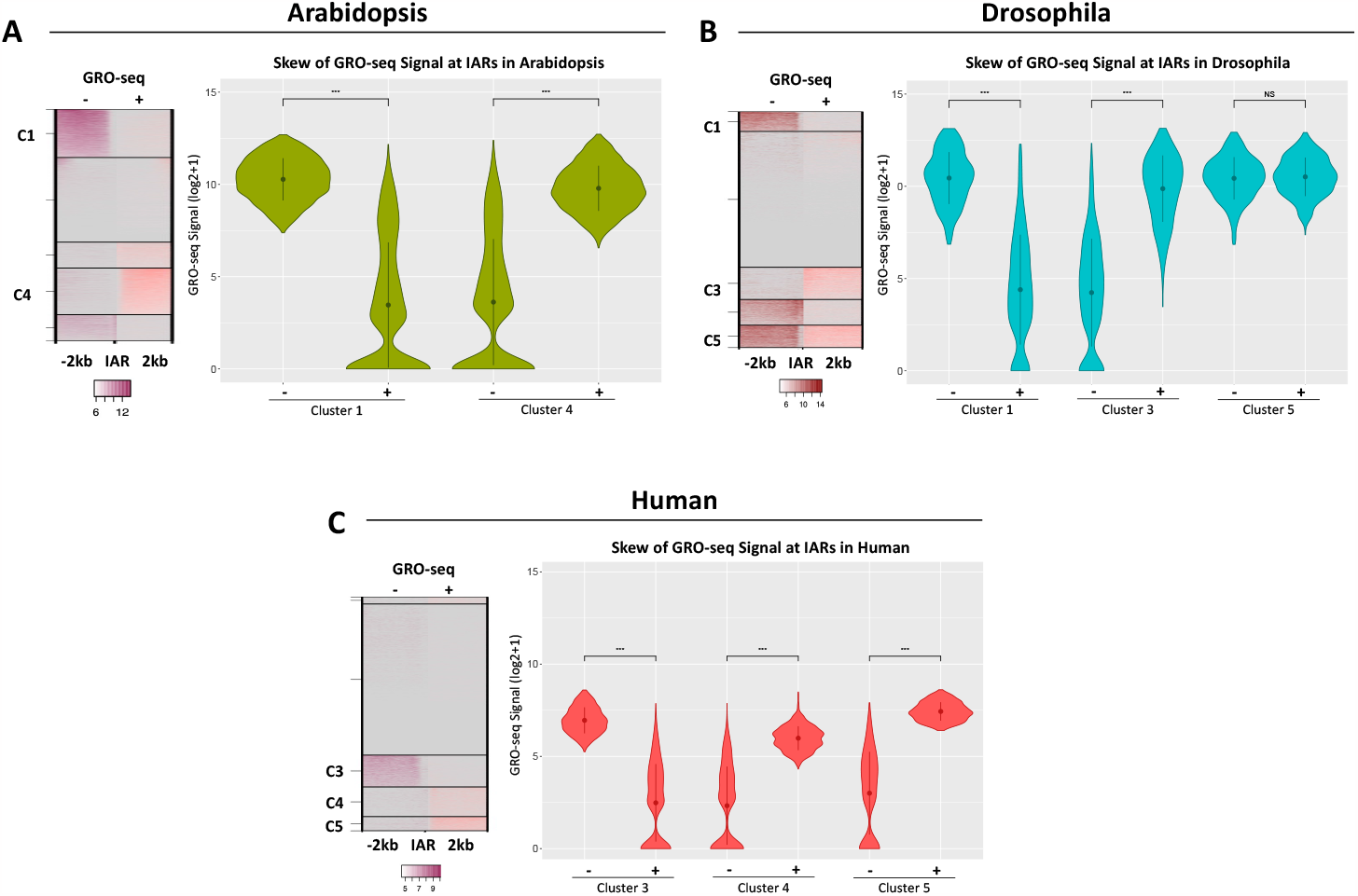
Skew of GRO-seq signal across intergenic accessible regions (IARs). GRO-seq signal was quantified on the upstream and downstream sides of each IAR to determine directionality of signal for **A)** Arabidopsis, **B)** Drosophila and **C)** Human. Directionality was calculated only for clusters that had strong GRO-seq signal on one or both sides of the IAR. Loci with low or no signal on either side were excluded from calculations. GRO-seq signal has been (log2+1) transformed. Statistical significance between up and downstream signal was calculated using a Wilcoxon Rank Sum test with a Bonferroni corrected significance threshold.

### ANALYSIS OF GENIC PTMs IN OTHER FLOWERING PLANT SPECIES SUGGESTS A SIMILAR UNIDIRECTIONAL TRANSCRIPTIONAL MECHANISM AT GENES

In order to assess whether unidirectional transcription initiation at genes is unique to Arabidopsis or a characteristic of flowering plants more generally, we analyzed publicly available chromatin accessibility and ChIP-seq data from *Oryza sativa* (rice) and *Glycine max* (soybean) (Lu *et al*. 2019). We chose rice and soybean for these analyses to encapsulate both long evolutionary distances as well as different genome sizes. In our comparisons of rice, a monocot with a relatively small genome, and soybean, a dicot with a larger genome, we examined chromatin marks around the TSSs of plus and minus strand genes as a proxy for transcriptional directionality, as we did not have access to nascent transcript datasets for these species. Consistent with our observations in Arabidopsis, we observe one-sided flanking of transcription-associated histone modifications H3K4me1 and H3K4me3, as well as H2A.Z in the direction of the gene body (**Figure 4**). Thus, these plants also show a pattern of PTMs consistent with unidirectional transcription at TSSs, suggesting that this may represent a general difference between metazoans and flowering plants. This is also consistent with nascent RNA analyses in maize and cassava, which found no evidence for extensive divergent transcription at genic TSSs (Erhard *et al*. 2015; Lozano *et al*. 2021).

**Figure 4:**
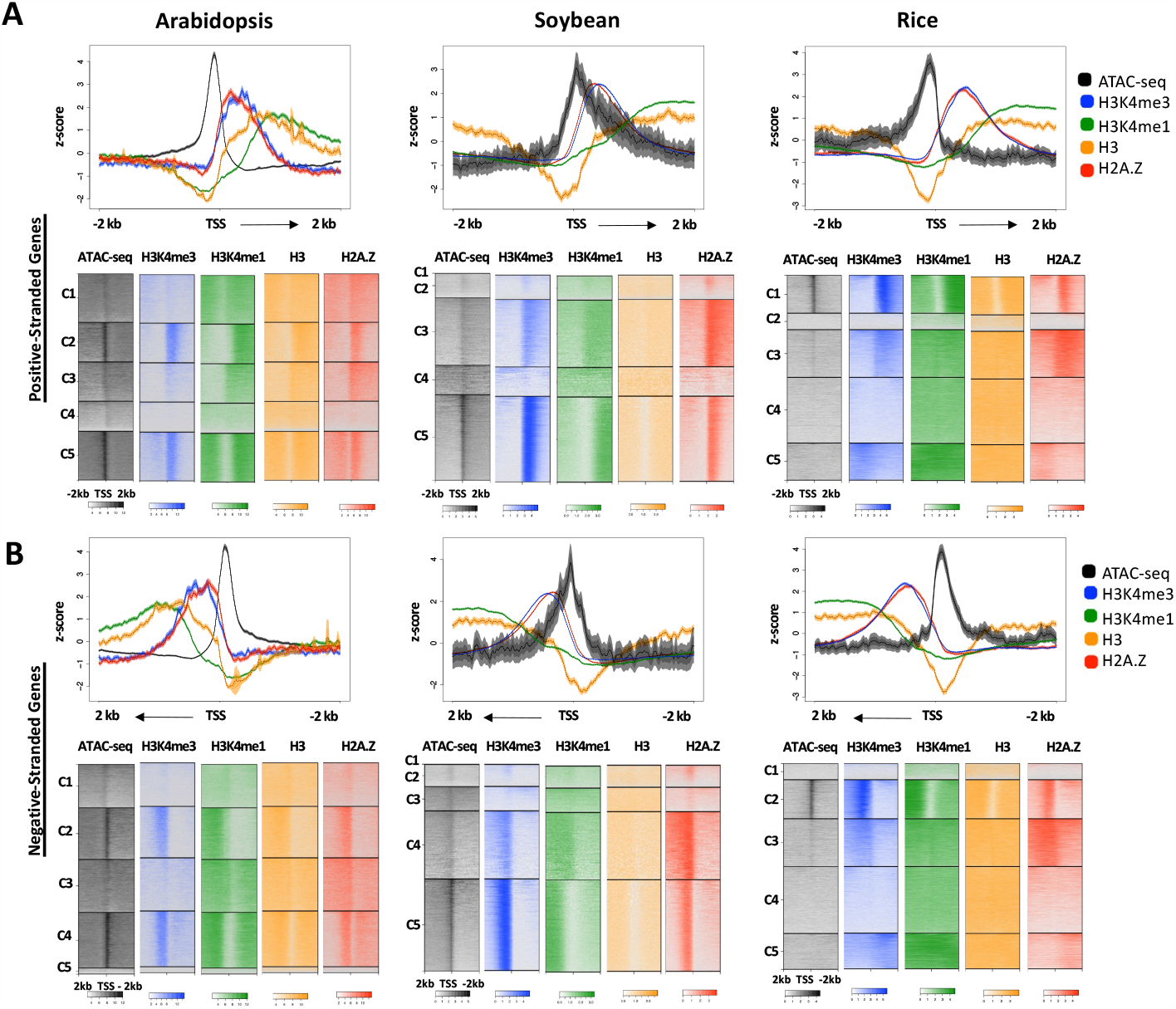
Enrichment patterns around the TSS in multiple angiosperm species. **A)** Metaplots and heatmaps of average H3K4me3, H3K4me1, H3 and H2A.Z signal and chromatin accessibility across positive-stranded gene TSSs in Arabidopsis (14,420 TSS sites), soybean (28,691 TSS sites), and rice (49,066 TSS sites). **B)** Metaplots and heatmaps of average H3K4me3 and H3K4me1, H3 and H2A.Z signal and chromatin accessibility across negative-stranded TSSs in Arabidopsis, soybean, and rice. Windows extend 2 kb upstream and 2 kb downstream of the TSS. In the metaplots, solid lines represent the mean signal intensity (z-score normalized); the inner, dark-shaded region represents the standard error; and the outer, light-shaded region represents the 95% confidence interval (CI) of signal intensity. Heatmaps have been log2 transformed.

## DISCUSSION

In this study, we confirm the unique unidirectional histone PTM enrichment observed previously in *Arabidopsis* genes and show that this feature correlates with the direction of nascent transcription, a feature also present at intergenic CREs. Our study also provides ChIP-seq datasets from the root non-hair cell type for the highly conserved set of histone PTMs H3K27ac, H3K27me3, H3K4me1, and H3K4me3, as well as histone variant H2A.Z in the model plant *Arabidopsis thaliana*. When examined alongside ATAC-seq data previously generated by our lab and available GRO-seq, NET-seq and 5-EU RNA-seq data, stark differences become apparent between plant and animal species. Histone PTM enrichment at TSSs of protein-coding genes in human and Drosophila show a distinct bimodal distribution pattern around the TSS, while the enrichment pattern found around TSSs in Arabidopsis and two other plant species is noticeably missing signal upstream of the TSS (**Figures 1 and 4**). This bimodal pattern has been shown to be indicative of bidirectional transcription at the promoter (Kapranov *et al*. 2007; Core *et al*. 2008; Seila *et al*. 2008), strongly implying that in Arabidopsis, rice, and soybean, transcription is more tightly regulated at the level of initiation in terms of directionality.

Proximal intergenic CREs in animal genomes correspond with the hallmark enhancer histone PTM pattern, being flanked by nucleosomes with high levels H3K27ac (active) or H3K27me3 (poised) and H3K4me1, with relatively less H3K4me3 **(Figure 2)**. While metaplots of PTMs at intergenic accessible sites suggested a similar bimodal pattern in Arabidopsis, clustered heatmaps revealed that histone PTMs flank the region either upstream or downstream of the accessible site. Additionally, H3K27me3 and H3K27ac do not appear to be mutually exclusive, as they are in animal species. Taken together, these findings indicate that the characteristic bimodal PTM distribution used in animals to identify enhancer regions will not apply to Arabidopsis and likely other plant species, where these marks generally flank only the single transcribed side of the element. Finally, nascent transcript data reveal that putative eRNAs at Arabidopsis CREs are produced mostly unidirectionally (**Figure 2C**), consistent with recent findings in Arabidopsis immunity-related CREs (Zhang *et al*. 2022). In contrast, a previous study of nascent transcription in cassava and maize using PRO-seq did report bidirectional transcription from many intergenic sites in both species (Lozano *et al*. 2021). Thus, the extent to which bidirectional CRE transcription is restricted in plant genomes, as well as the gene structure and/or promoter:enhancer interactions that differ between CREs that produce unidirectional eRNAs and those that are bidirectional will be rich areas for future study.

Overall, the results of this investigation indicate that genuine differences exist between the plant and animal kingdom at the level of transcriptional initiation. While the elongation of protein-coding transcripts in the sense direction appears to be preferred across all eukaryotes, the results of this study support the notion that this direction is preferred with near exclusivity in transcriptional initiation at TSSs in plants, while animal transcription initiation is more promiscuous.

These results beg the question, what is the reason for this stark contrast in transcriptional directionality between plant and animal species? One potential reason is that, distinct from animals, plants contain RNA-directed DNA methylation pathways that can silence portions of the genome, and these pathways are primarily targeted by siRNAs, which could be generated by overlapping divergent transcripts. If allowed to proceed unchecked, the production of such reverse transcripts could disrupt the plant epigenome, resulting in a strong selective pressure to keep the generation of bidirectional RNAs tightly regulated.

How, then, are plants able to prevent the production of reverse strand transcripts? Recent findings from Hi-C data with single-gene resolution may shed some light on the question. Rather than forming the large topologically associated domains (TADs) found in mammals, the *Arabidopsis* genome is preferentially organized into small, local gene loops, where the 5’ and 3’ ends of a gene directly interact (Liu *et al*. 2016). This organization was also recently reported in maize, tomato, and the liverwort *Marchantia polymorpha*, suggesting that the gene loop is a common organizational feature of plant genomes (Lee and Seo 2023). The constrained geometry of these gene loops has been suggested to eliminate bidirectional transcription, forcing Pol II to transcribe in the sense direction alone (Tan-Wong *et al*. 2012). This organizational scheme and relative lack of long-range interactions could explain why the majority of intergenic accessible chromatin sites we identified were preferentially located proximally upstream of their nearest gene (Maher *et al*. 2018). The Pol II-associated factor Ssu72 is responsible for maintaining the association between the gene ends in yeast; when this factor is mutated, the gene loop structure is abolished and bidirectional transcripts are produced (Tan-Wong *et al*. 2012; Castelnuovo and Stutz 2013). While it is not yet known whether plants contain a functional counterpart of Ssu72, these findings suggest that analogous differences in higher order chromatin structure may be responsible for the observed distinctions in transcriptional directionality.

It is clear that many histone PTMs are deposited as the result of active transcription and are strongly correlated with transcriptional activity because they are the effect, not the cause, and instead provide a supportive role for continual transcription. However, targeted reversal of transcription-supportive chromatin modifications may still play a role in regulating transcriptional directionality. For example, the histone demethylase Flowering Locus D has been found to help limit transcription in regions of convergent genes to prevent accumulation of antisense RNAs by limiting the abundance of H3K4me1 (Inagaki *et al*. 2021). Similarly, studies in yeast have shown that the Hda1 histone deacetylase complex represses divergent transcription by deacetylating histone H3 (Gowthaman *et al*. 2021).

The presence of histone variants such as H2A.Z can also play a role in supporting transcription, and thus may also be a point of control for regulating transcriptional initiation directionality. For example, H2A.Z can promote transcription by lowering the barrier for Pol II progression (Weber *et al*. 2014). In animals, H2A.Z is often found at well-positioned -1 and +1 nucleosomes flanking the TSS of actively transcribed genes, while in Arabidopsis and other plants, it is generally only found at the +1 nucleosome. Therefore, another possibility is that targeted removal or inhibition of deposition of histone variants such as H2A.Z could also modulate the direction in which transcription is likely to initiate. The interplay between gene organization, higher order genome structure, and targeted chromatin modifications likely all contribute to regulating transcription initiation directionality and will be fruitful to consider together in future studies on this topic.

## MATERIALS and METHODS

### Publicly available datasets

Publicly accessible datasets from the Encyclopedia of DNA Elements (ENCODE) project (2004, 2012) (https://www.encodeproject.org/) and the Gene Expression Omnibus (GEO) (Edgar *et al*. 2002) (https://www.ncbi.nlm.nih.gov/geo/) were used in this study. Details about each of these datasets, including the species and cell type/tissue used, the accession numbers for each library, and the genome version that these data were mapped to in our study, are detailed in **Supplementary Table 1**.

### Preparation of Arabidopsis ChIP-seq libraries

Non-hair cell nuclei were isolated from *A. thaliana* (Col-0) roots carrying the GL2p:NTF and ACT2p:BirA transgenes using the INTACT method as described previously (Wang and Deal 2015). ChIP-seq libraries were prepared and sequenced as described in(Adli and Bernstein 2011). The antibodies used to prepare the ChIP-seq libraries are listed in **Supplementary Table 2**.

### Data analysis

Raw sequence read processing, mapping, peak calling, and genomic distribution determination were all conducted as described previously (Maher *et al*. 2018). **Supplementary Table 3** details the data quality of the *A. thaliana* non-hair cell ChIP-seq generated by this study. After processing and mapping, GRO-seq data were run through an R script which removed the top 10% of reads to prevent high-signal artifacts from skewing the distribution of the metaplots. Heatmaps and metaplots were generated using SeqPlots (http://seqplots.ga/) (Stempor and Ahringer 2016).

### Calculation of Skew

Upstream and downstream bedfiles were generated by calculating the midpoint of the intergenic accessible sites from the original bedfile and then adding/subtracting 2 kb. Bigwig signal from all ChIP and GRO-seq datasets were obtained using bigWigAverageOverBed (https://www.encodeproject.org/software/bigwigaverageoverbed/) from kentUtils (Kent *et al*. 2010). Results were filtered to ignore data points with no signal on either side of the IAR. Statistical significance was determined using a Wilcoxon Rank Sum Test to determine whether there were significant differences between (log2+1) transformed signal intensity for upstream and downstream regions, with a Bonferroni-corrected threshold applied to each of these (Curtin and Schulz 1998). Finally, data was visualized via violin plot (ggplot2) (Wickham 2016).

### Data availability

ChIP-seq data from Arabidopsis root non-hair cells are deposited in the NCBI GEO database under accession number GSE152243. All other datasets are previously published and their accession numbers can be found in Supplementary Table 1.

## ACKNOWLEDGEMENTS

We would like to thank the members of the Deal Lab for their feedback and support. We would also like to extend a special thanks to Benjamin Barwick, Ph.D., Marko Bajic, Ph.D. and David Gorkin, Ph.D. for their advice regarding data analyses.

## FUNDING

This work was supported by funding from Emory University and the National Institutes of Health (R01GM134245) to RBD. BDS was also supported by an NIH training grant (T32GM008490).

## SUPPLEMENTARY FIGURES

**Supplementary Figure 1:**
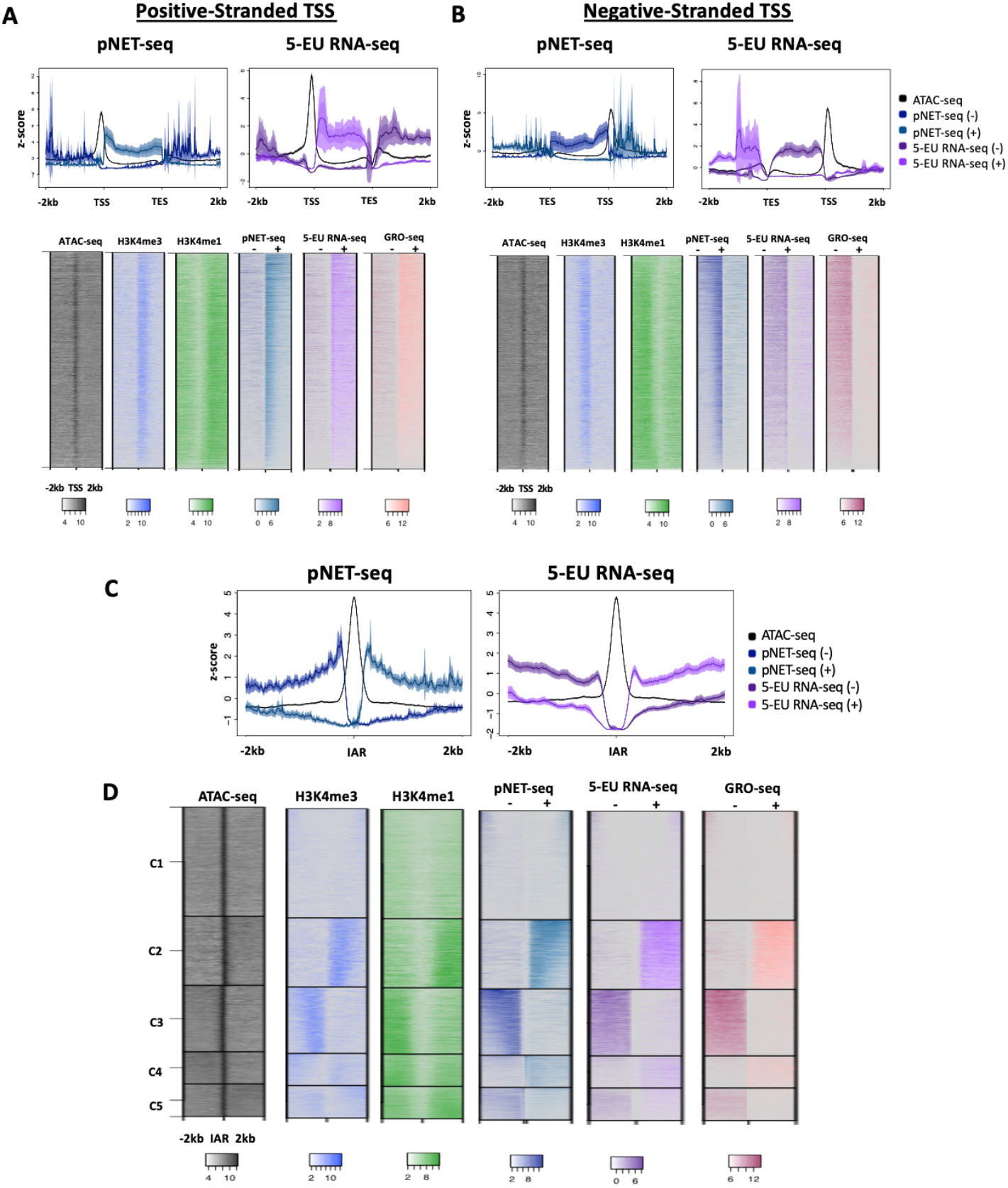
Additional nascent RNA-seq datasets at TSSs and IARs in Arabidopsis thaliana. 5-EU RNA-seq and pNET-seq are shown as average plots over gene bodies (top images) at Arabidopsis positive-stranded TSSs (**A**) as well as negative-stranded TSSs (**B**). Solid lines represent the mean signal intensity (z-score transformed); the inner, dark-shaded region represents the standard error; and the outer, light-shaded region represents the 95% confidence interval (CI) of signal intensity. Lower images in each panel show the nascent RNA data in heatmap form, along with ChIP-seq data for H3K4me1 and H3K4me3. (**C**). Average plots of nascent RNA signals at Arabidopsis intergenic accessible chromatin regions (accessible sites outside of gene bodies and 100-2000 bp upstream of a TSS). (**D**) Images in each panel are K-means clustered heatmaps containing all three types of nascent RNA data, along with ChIP-seq data for H3K4me1 and H3K4me3 at intergenic accessible regions.

**Supplementary Figure 2:**
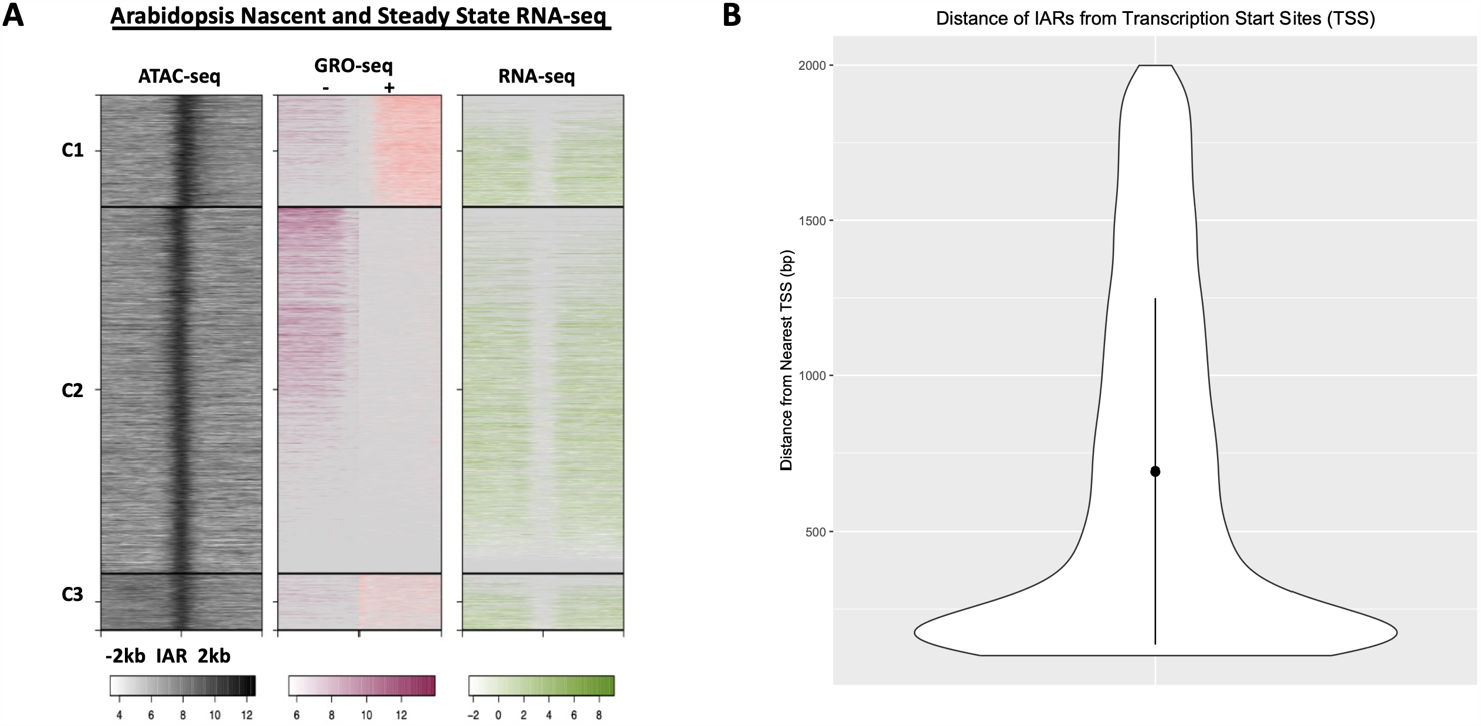
Comparison of nascent and steady-state RNA-seq at intergenic enhancer regions in Arabidopsis. **A)** Clustered heatmaps of ATAC-seq, GRO-seq, and RNA-seq at intergenic accessible regions (IARs), which are defined as accessible chromatin sites outside of transcribed protein coding genes and within the range of 100-2000 bp from the nearest TSS. Windows extend 2 kb upstream and 2 kb downstream of the IARs; heatmaps have been log2 transformed. **B)** Distribution of distances between IARs and TSSs, showing an average distance of 692 bp from a TSS.

## SUPPLEMENTARY TABLES

**Supplementary Table 1:**
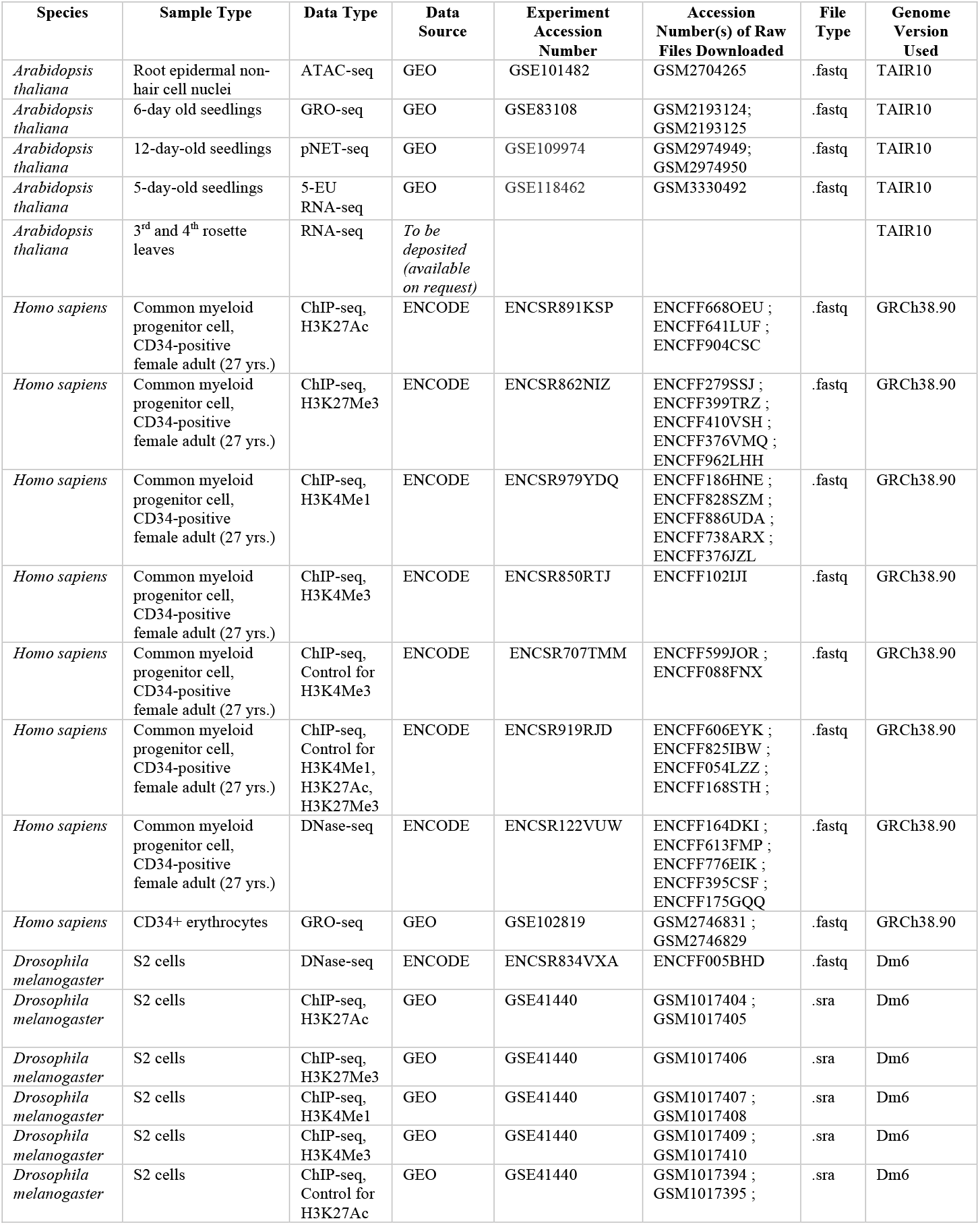
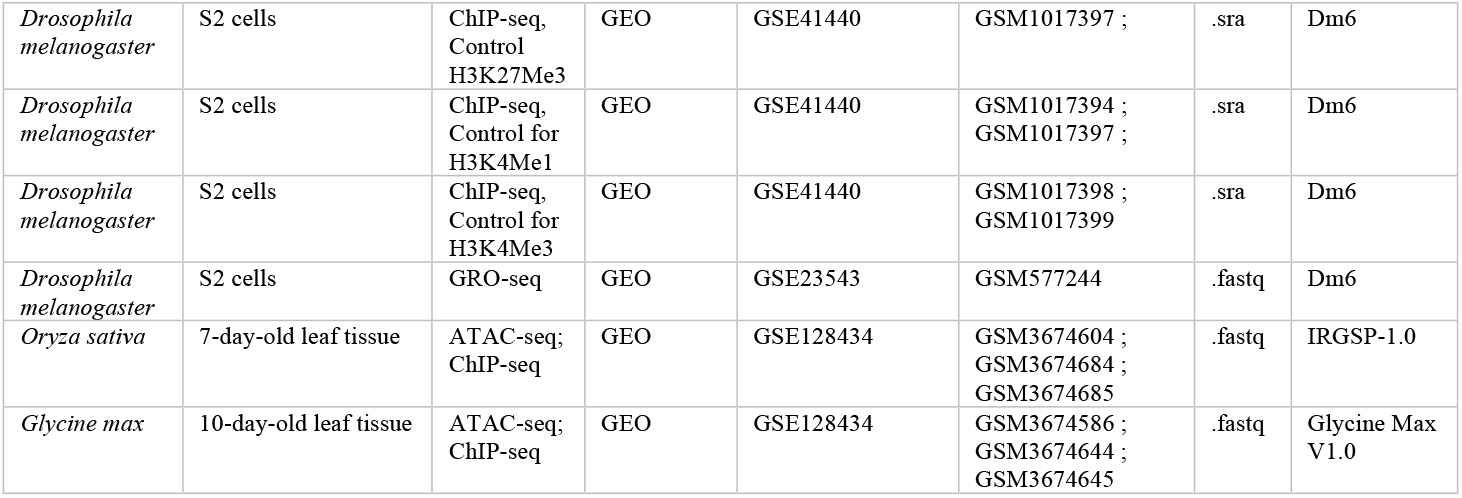
Publicly available dataset information.

**Supplementary Table 2:** *A. thaliana* ChIP-seq antibody information.

| Target | Antibody Name | Supplier | Concentration | Quantity Used per Reaction |
| --- | --- | --- | --- | --- |
| H3K4me1 | ab8895 | Abcam | 0.5 mg/mL | 2 µg |
| H3K4me3 | ab8580 | Abcam | 0.45 mg/mL | 1.8 µg |
| H3K27ac | ab4729 | Abcam | 0.5 mg/mL | 2 µg |
| H3K27me3 | 07-449 | Millipore | 0.5 mg/mL | 2 µg |
| H3 | ab1791 | Abcam | 0.5 mg/mL | 2 µg |

**Supplementary Table 3:** Data quality of *A. thaliana* root epidermal non-hair cell ChIP-seq datasets.

| Dataset type | Read size (nt) | Single end (SE) or paired end (PE) | Total reads | Total mapped reads | Total mapped q2 filtered reads | Total nuclear peaks called (via HOMER) | Avg. size of peaks (bp) | Std. dev. of peak size (+/- bp) | Median size of peaks (bp) |
| --- | --- | --- | --- | --- | --- | --- | --- | --- | --- |
|  |  |  | (x 10<sup>6</sup>) | (x 10<sup>6</sup>) | (x 10<sup>6</sup>) |  |  |  |  |
| ChIP-seq (H3K4me1) | 50 | SE | 83.2 | 73.1 | 54.7 | 31,016 | 402.54 | 201.26 | 330 |
| ChIP-seq (H3K4me3) | 50 | SE | 101.1 | 91.5 | 82.6 | 14,718 | 277.12 | 134.57 | 189 |
| ChIP-seq (H3K27ac) | 50 | SE | 131.6 | 115 | 92 | 36,146 | 299.09 | 161.41 | 235 |
| ChIP-seq (H3K27me3) | 50 | SE | 22.5 | 19.8 | 16.1 | 27,784 | 378.9 | 223.91 | 303 |
| ChIP-seq (H2A.Z) | 50 | SE | 123.3 | 104.6 | 81.8 | 30,594 | 255.62 | 115.61 | 183 |
| ChIP-seq (H3) | 50 | SE | 62 | 53.8 | 34.3 | 23,379 | 254.38 | 76.43 | 211 |

**Supplementary Table 4.** Summary statistics for raw GRO-seq signal enrichment up and downstream of IARs for each species. Resulting un-transformed values of GRO-seq signal enrichment for each cluster, corresponding to Figure 2.

| Arabidopsis: Downstream |  |  |  | Arabidopsis: Upstream |  |  |  |
| --- | --- | --- | --- | --- | --- | --- | --- |
| Cluster ID | Mean | Median | Standard Dev | Cluster ID | Mean | Median | Standard Dev |
| 1 | 129 | 5.44 | 328 | 1 | 1661 | 1259 | 1272 |
| 2 | 65.6 | 5.44 | 267 | 2 | 84.4 | 8.74 | 311 |
| 3 | 416 | 151 | 678 | 3 | 74.9 | 8.74 | 241 |
| 4 | 1257 | 859 | 1133 | 4 | 145 | 8.74 | 387 |
| 5 | 63.2 | 5.44 | 208 | 5 | 543 | 204 | 785 |
| Drosophila: Downstream |  |  |  | Drosophila: Upstream |  |  |  |
| Cluster ID | Mean | Median | Standard Dev | Cluster ID | Mean | Median | Standard Dev |
| 1 | 173 | 19.6 | 572 | 1 | 2143 | 1451 | 1983 |
| 2 | 131 | 0 | 665 | 2 | 59.5 | 0 | 397 |
| 3 | 1721 | 1126 | 1769 | 3 | 120 | 19.1 | 359 |
| 4 | 173 | 30 | 404 | 4 | 1277 | 703 | 1467 |
| 5 | 1870 | 1467 | 1348 | 5 | 1844 | 1475 | 1441 |
| Human: Downstream |  |  |  | Human: Upstream |  |  |  |
| Cluster ID | Mean | Median | Standard Dev | Cluster ID | Mean | Median | Standard Dev |
| 1 | 40.1 | 36.6 | 32.5 | 1 | 9.7 | 0 | 20 |
| 2 | 4.44 | 0 | 12.4 | 2 | 8.24 | 0 | 18.4 |
| 3 | 13.4 | 4.65 | 22 | 3 | 139 | 119 | 71.1 |
| 4 | 69.3 | 61.5 | 35.1 | 4 | 12.1 | 5.62 | 19.2 |
| 5 | 183 | 167 | 65.5 | 5 | 19.9 | 11.2 | 28.6 |

## Notes

### Competing Interest Statement

The authors have declared no competing interest.

